# Persistent Inward Currents Increase With The Level Of Voluntary Drive In Plantar Flexor Low-Threshold Motor Units

**DOI:** 10.1101/2020.12.11.420570

**Authors:** Lucas B. R. Orssatto, Karen Mackay, Anthony J. Shield, Raphael L. Sakugawa, Anthony J. Blazevich, Gabriel S. Trajano

## Abstract

This study tested the hypothesis that estimates of persistent inward currents (PICs) in the human plantar flexors would increase with the level of voluntary drive. Twenty-one participants volunteered for this study (29.2±2.6 years). High-density surface electromyograms were collected from *soleus* and *gastrocnemius medialis* during ramp-shaped isometric contractions to 10%, 20%, and 30% (torque rise of 2%/s and 30-s duration) of each participant’s maximal torque. Motor units identified in all the contraction intensities were included in the paired-motor unit analysis to calculate delta frequency (ΔF) and estimate the PICs. Increases in PICs were observed from 10% to 20% (Δ=0.6 pps; p<0.001) and 20% to 30% (Δ=0.5 pps; p<0.001) in *soleus*, and from 10% to 20% (Δ=1.2 pps; p<0.001) but not 20% to 30% (Δ=0.09 pps; p=0.724) in *gastrocnemius medialis*. Maximal discharge rate increased for *soleus* and *gastrocnemius medialis* from 10% to 20% (respectively, Δ=1.75 pps, p<0.001; and Δ=2.43 pps, p<0.001) and 20% to 30% (respectively, Δ=0.80 pps, p<0.017; and Δ=0.92 pps, p=002). The repeated-measures correlation identified associations between ΔF and increases in maximal discharge rate for both *soleus* (r=0.64; p<0.001) and *gastrocnemius medialis* (r=0.77; p<0.001). An increase in voluntary drive tends to increase PIC strength, which has key implications for the control of force but also for comparisons between muscles or between studies when relative force levels might be different. These data indicate that increases in voluntary descending drive amplify PICs in humans and provide an important spinal mechanism for motor unit firing, and thus force output modulation.

## INTRODUCTION

The force produced by a skeletal muscle during contraction is largely dependent upon the number of recruited motor units and their discharge rates (Enoka and Duchateau, 2017). In the presence of monoaminergic input (i.e. neuromodulation), motor neurone discharge rates are increased by the development of persistent inward currents (PICs) (Hounsgaard *et al*., 1988; Lee and Heckman, 1999, 2000), which are depolarising currents generated by voltage-sensitive sodium and calcium channels primarily located on motor neurones (Eckert and Lux, 1976; Schwindt and Crill, 1977; Hounsgaard and Kiehn, 1993). PICs amplify and prolong excitatory synaptic input, allowing a greater muscle force production for a given level of descending drive as well as the capacity to maintain contraction even as descending drive is reduced below the level required for initial muscle activation. Monoamines such as serotonin and noradrenaline facilitate PICs, changing the input-output gain of motor neurones up to 5-fold (Hounsgaard *et al*., 1988; Lee and Heckman, 1999, 2000). Additionally, the motor system would benefit from gain adjustments in order to produce varying levels of force, and it has been hypothesised that PICs could act as a variable gain control system by introducing non-linearities in the input-output relationship, offering an elegant solution to appropriately amplifying inputs to achieve forces across a diverse range of motor activities (Johnson and Heckman, 2014). Both animal experiments (Powers, Nardelli and Cope, 2008; Huh *et al*., 2017) and computational models (Heckman and Binder, 1991; Heckman, 1994; Randall K. Powers and Heckman, 2015) indicate that a non-linear relationship exists between the ionotropic input and the excitatory motor output, which depends on the intensity of the muscle contraction (Naufel *et al*., 2019). Thus, PICs may act as the main gain control mechanism within motor neurones (Johnson and Heckman, 2014; Binder, Powers and Heckman, 2020), playing an important role in normal motor behaviour by allowing optimum muscle force control.

In human motor neurones it is not possible to directly measure PIC amplitudes. However, by using the discharge rates of motor units during voluntary contractions, PIC amplitudes can be estimated using the paired motor unit technique (Gorassini *et al*., 2002; Powers, Nardelli and Cope, 2008; Stephenson and Maluf, 2011). Paired motor unit analysis consists of pairing the discharge rates of a low threshold or *control unit* to a higher threshold or *test unit* (Gorassini *et al*., 2002; Powers, Nardelli and Cope, 2008;

Stephenson and Maluf, 2011). These are obtained during a slowly-increasing and - decreasing ramped contraction. The difference in discharge rate of the control unit at the time of recruitment and de-recruitment of the test unit is calculated and referred to as the Δ frequency (ΔF). ΔF has been validated using both animal and computer models, and is considered to be proportional to PIC amplitude (Powers, Nardelli and Cope, 2008; Randall K Powers and Heckman, 2015). Therefore, a higher ΔF, representing a greater difference in recruitment and de-recruitment discharge rates, indicates a higher PIC amplitude.

Contraction intensity-dependent PIC amplification has significant implications for our understanding of the modulation of skeletal muscle force. However, despite the increasing evidence of intensity-dependent PIC amplification, it is still not known whether this phenomenon exists in human motor neurones. Although the extrapolation of gain-control physiology from animal preparations or computational simulations to human physiology are comprehensible, it has been shown that PIC amplification varies between animals and within muscle groups (Heckman, Gorassini and Bennett, 2005; Heckman *et al*., 2009; Heckman and Enoka, 2012; Huh *et al*., 2017; Manuel *et al*., 2019; Binder, Powers and Heckman, 2020), so direct testing across human muscles is needed (Afsharipour *et al*., 2020; Kim *et al*., 2020). In fact, two recent studies (Afsharipour *et al*., 2020; Kim *et al*., 2020) reported no differences in the PIC amplitudes (estimated through paired-motor unit ΔF analysis) across different motor outputs (10%, 20%, and 30% of maximal voluntary force). Nevertheless, these results should be interpreted cautiously because of their methodological design. They adopted a different rate of torque rise and decline among intensities, which could have underestimated the ΔFs obtained at higher intensities due to spike threshold accommodation (Randall K. Powers and Heckman, 2015). Also, they did not compare the same motor units across contraction intensities. These studies used surface high-density electromyography and algorithm decomposition to investigate the discharge patterns of motor units across different intensities. However, because the decomposition algorithm tends to identify a lower proportion of lower-threshold motor units during higher intensities contractions (Hassan *et al*., 2019), they might have compared different motor units with distinct recruitment characteristics. Aiming to address the limitations of the previous studies, we tested the hypothesis that PIC amplitudes would increase according to muscle contraction intensity in human *soleus* and *gastrocnemius medialis* adopting the same rate of torque rise and contraction duration, and tracking the same motor units across different contraction intensities.

## MATERIALS AND METHODS

### Participants and ethical procedures

To participate, volunteers had to be free from neural and musculoskeletal disorders in the lower limb. All participants were asked to avoid coffee and medications that could influence monoaminergic release, such as serotonin or noradrenaline. Furthermore, they were asked to abstain from exercising 24 h prior to the testing session. Participants were excluded from the data analyses if: a) no usable motor units were identified by the decomposition algorithm; or b) if it was not possible to pair available motor units (see details below). This study was approved by the local Human Research Ethical Committee of the Queensland University of Technology, and all participants gave written informed consent prior to participation in the study.

### Study design and neuromuscular testing procedures

Participants visited the laboratory on one occasion, during which participants were familiarised with testing procedures and data were collected from the plantar flexors. After electrode placement on *soleus* and *gastrocnemius medialis*, the participants were seated upright on the chair of an isokinetic dynamometer (Biodex System 4, Biodex Medical system, Shirley, NY) with the knee fully extended (0°) and ankle at neutral position (0°). A warm-up consisting of six 5-s voluntary submaximal isometric plantar flexion contractions (2 × 40%, 2 × 60% and 2 × 80% of their perceived maximal effort) was performed, followed by three maximal voluntary contractions of ~3-s with 30-s rest intervals. The maximum torque achieved was recorded as their maximal voluntary torque. Next, participants were familiarised with both trapezoidal and triangular ramped contractions. Both triangular and trapezoidal contractions have been used previously to calculate ΔFs using the paired motor unit analysis (Wilson *et al*., 2015; Hassan *et al*., 2020). These ramp-shaped contractions were performed to 10, 20 and 30% of their maximal voluntary contraction torque. All contractions had a duration of 30 s and a rate of torque increase and decrease of 2%/s. Contraction duration was equalised between tests at different intensities because longer muscle contractions result in a spike frequency adaptation, making motor units discharge at lower frequencies for a given force, inflating the ΔFs (Vandenberk and Kalmar, 2014). Also, rate of torque production was identical between contraction intensities because faster rates can reduce ΔFs due to spike-threshold accommodation (Vandenberk and Kalmar, 2014). Therefore to control the torque condition in which we compared the three intensities, the same amount of torque (10% of MVC) with the same rate of torque rise (2% per s) was observed in the initial and final 5 s of each contraction intensity. Participants were instructed to follow a torque path provided in real time on a large computer monitor (Figure 1) during each ramp-shaped contraction. Five minutes after the familiarisation, the order of the ramped contractions was randomised, and data collection proceeded. Participants performed 2-3 attempts at each intensity with 40-s rest intervals. When an abrupt increase or decrease in torque was observed, the respective contraction was excluded.

**Figure 1.**
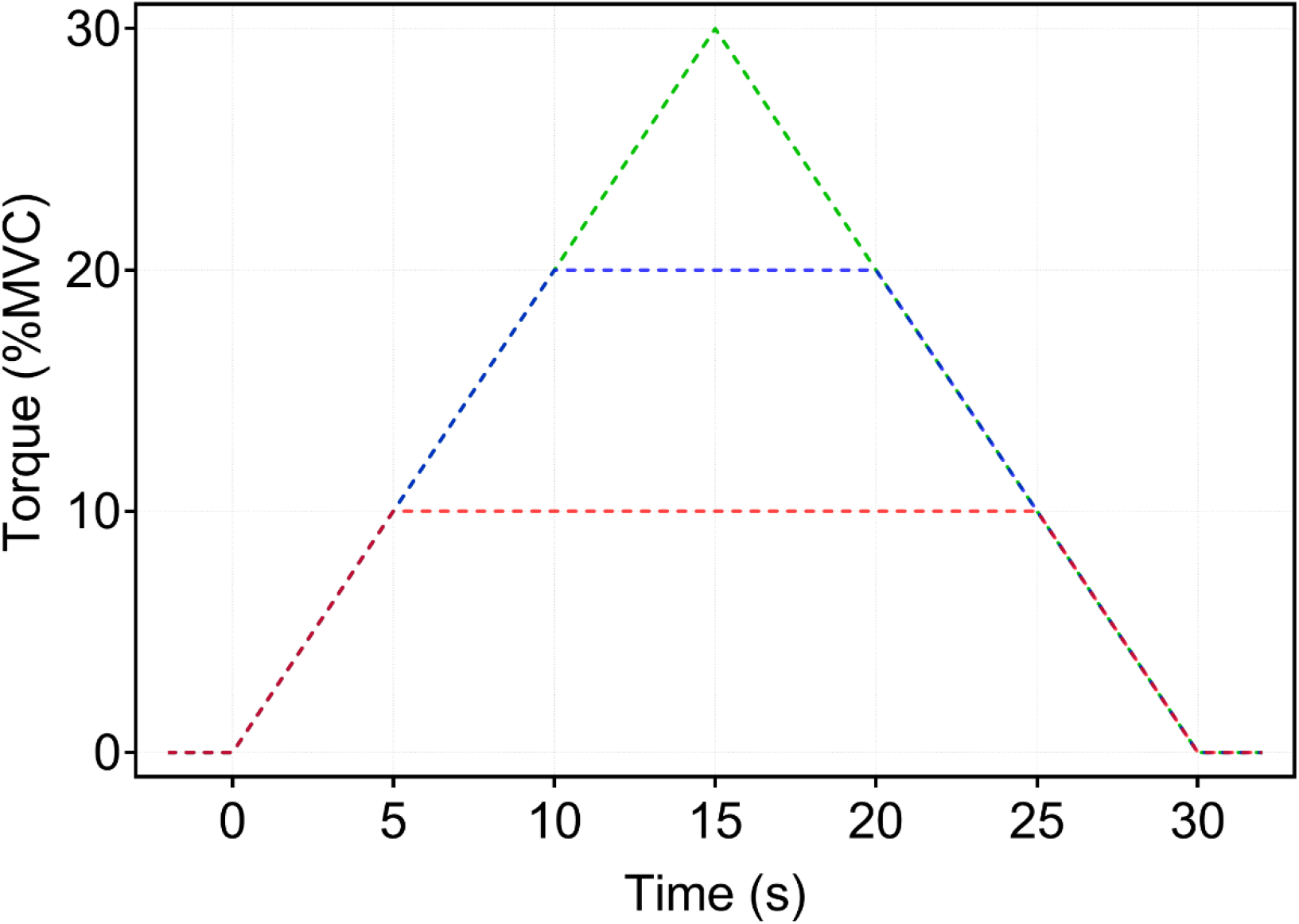
Torque path for ramp-shaped contractions at each contraction intensity. Required torque path for the ramp-shaped contractions with 10% (red), 20% (blue) and 30% (green) of the participant’s maximal voluntary torque. A torque rise rate of 2%/s was adopted for all contractions. For the 10% and 20% contractions, 20-s and 10-s sustained torque contractions phases were included, respectively, to ensure that all contractions lasted 30 s.

### Surface electromyography (sEMG)

sEMG was recorded during submaximal ramped contractions using two semi-disposable 32-channel electrode grids with a 10-mm inter-electrode distance (ELSCH032NM6, OTBioelettronica, Torino, Italy). After skin shaving, abrasion, and cleansing with 70% isopropyl alcohol, electrodes were placed over the muscle belly of *soleus* and *gastrocnemius medialis*, on the medial portion of each muscle angled slightly towards the calcaneus tendon, using a bi-adhesive foam layer and conductive paste (SpesMedica, Battipaglia, Italy). A strap electrode (WS2, OTBioelettronica, Torino, Italy) was dampened and positioned around the ankle joint as a ground electrode. The sEMG signals were recorded in monopolar mode and converted to digital signal by a 16-bit wireless amplifier (Sessantaquattro, OTBioelettronica, Torino, Italy) using OTBioLab+ software (version 1.3.0., OTBioelettronica, Torino, Italy). sEMG signals were amplified (256×), sampled at 2000 Hz and band pass filtered (10-500 Hz) before being stored for offline analysis.

### Data analysis

#### Motor unit identification and tracking

The recorded data were processed offline using the DEMUSE software (Holobar and Zazula, 2007). For each contraction intensity, only the ramp-shaped contraction yielding the lowest deviation from the torque trajectory was analysed. High-density sEMG signal were band-pass filtered (20-500 Hz) with a second-order, zero-lag Butterworth filter. Thereafter, the blind-source, convolutive kernel compensation method of separation was used for signal decomposition (Holobar and Zazula, 2007; Holobar, Minetto and Farina, 2014). To identify the same motor unit at each contraction intensity (i.e., 10, 20 and 30% of MVC), motor units were tracked using the motor unit filters methodology described in (Francic and Holobar, 2021). For this purpose, the recordings of different contraction levels were concatenated, and motor unit filters identified from individual contraction level applied to recordings of other two contraction levels, identifying the motor unit firing patterns across all the contraction levels. After removing the duplicates of motor units identified from two or more contraction levels, a trained investigator manually inspected and edited the discharge patterns of the motor units. Only motor units with a local pulse-to-noise ratio equal to or greater than 30 dB and presenting a physiological discharge pattern (visually inspected by an experienced researcher) were kept for further analysis (Holobar, Minetto and Farina, 2014).

#### Estimation of PIC amplitude (ΔF) and maximal discharge rate

The observed discharge events for each motor unit were converted into instantaneous discharge frequencies and fitted into a 5^th^ order polynomial function. The maximum value obtained from the polynomial curve was considered the maximal discharge rate. Thereafter, PIC amplitude was estimated using the paired motor unit analysis. Motor units with a low recruitment threshold (i.e., control units) were paired with higher recruitment threshold motor units (i.e., test units). ΔF was calculated as the change in discharge frequencies of the control motor unit from the moment of recruitment to the moment of de-recruitment of the test unit (Gorassini *et al*., 2002; Heckman, Gorassini and Bennett, 2005). In order to pair motor units, the following criteria were adopted: 1) rate-to-rate correlations between the smoothed discharge rate polynomials of the test and control units was r ≥ 0.7; 2) test units were recruited at least 1.0 s after the control units; and 3) the control unit did not show discharge rate saturation after the moment of test unit recruitment (> 0.5 pps) (Gorassini *et al*., 2002; Udina *et al*., 2010; Vandenberk and Kalmar, 2014; Binder, Powers and Heckman, 2020; Hassan *et al*., 2020). Furthermore, ΔFs were calculated only for pairs identified at all three contraction intensities, therefore including only motor units recruited from 0 to 10% of MVC, which were averaged to obtain a single ΔF for each motor unit. Subsequently, ΔFs and maximal discharge rates from all motor units were averaged to provide a single ΔF per participant per intensity. Figure 2 shows one example of the paired motor unit analysis for the same pairs of motor units identified at different contraction intensities.

**Figure 2.**
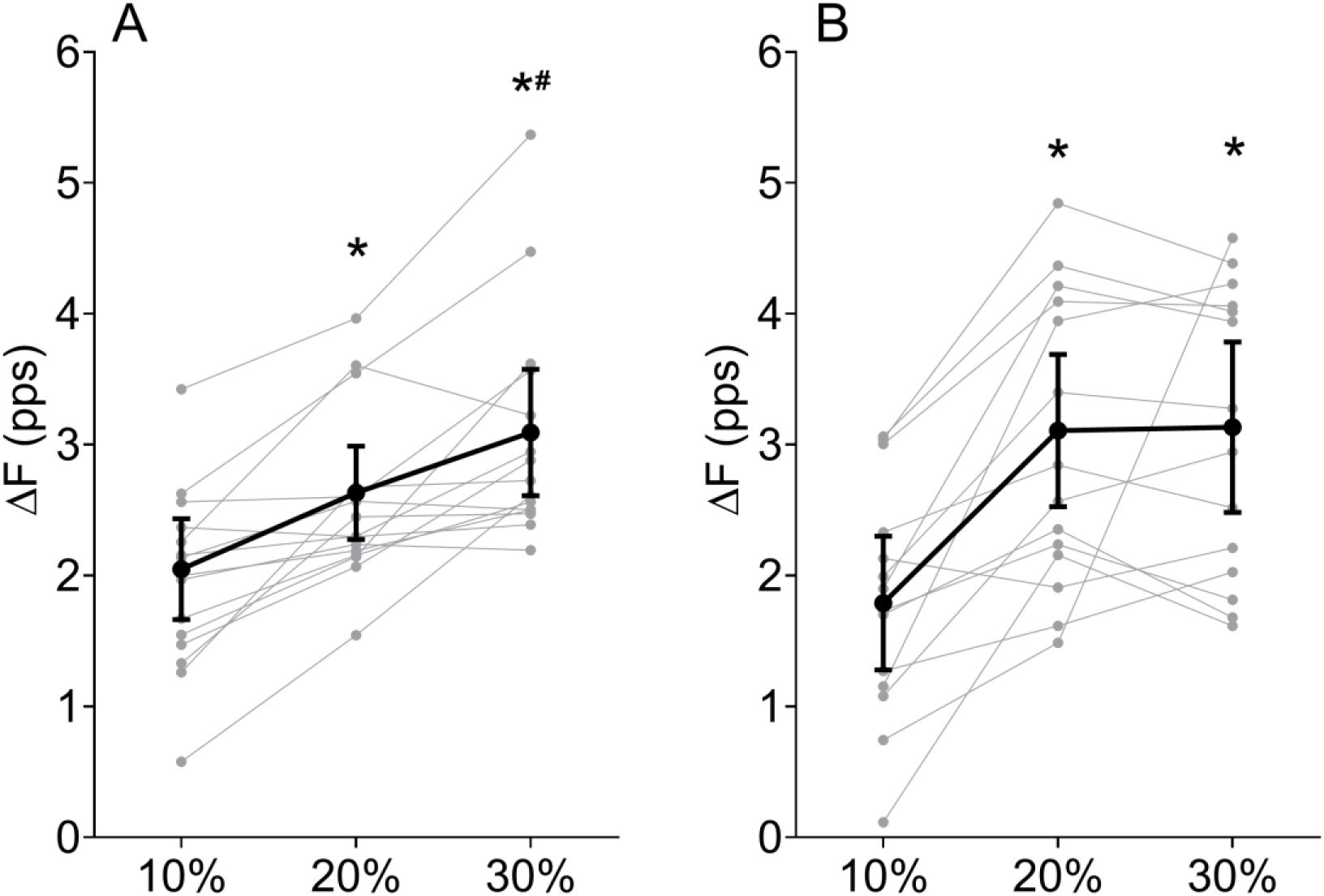
Delta frequency (ΔF) changes across different contraction intensities. ΔFs obtained in *soleus* (A) and gastrocnemius medialis (B) at 10%, 20% and 30% of maximal voluntary contraction. Grey dots and lines depict individual results (*soleus*, n = 15; *gastrocnemius medialis*, n = 14) and black dots and lines represent data mean ± 95% confidence interval lower and upper limits. * Different to 10% and ^#^ different to 20% conditions.

### Statistical analysis

A one-way repeated measure analysis of variance (ANOVA) was used to compare ΔFs and maximal discharge rates among different contraction intensities (10, 20 and 30%). If a significant effect was found, pairwise comparisons were performed using the Holm-Bonferroni post-hoc test. Equality of variance and sphericity assumptions were met for all tested variables. Therefore, it was not necessary to apply further corrections. The effect sizes derived from the ANOVA F ratios were calculated using the omega squared (ω^2^) method (0 - 0.01, very small; 0.01 – 0.06, small; 0.06 – 0.14, moderate; and > 0.14, large) (Lakens, 2013). In addition, the Cohen’s *d* effect size (presented as *d)* was calculated for pairwise comparisons (Lakens, 2013). All statistical tests were conducted using JASP software (version 0.11.1, University of Amsterdam, Netherlands) (JASP, 2019). Repeated-measures Bland–Altman within-subject correlation coefficients were subsequently computed using RStudio (Version 1.3.1056) and package “rmcorr” (Bakdash and Marusich, 2017). This method was used to determine the association between ΔFs and maximal discharge rates across the three contraction intensities (Bakdash and Marusich, 2017). Correlation magnitude was interpreted based on Cohen’s (1988) criteria: trivial, r < 0.1; small, r = 0.1–0.3; moderate, r = 0.3–0.5; large, r = 0.5–0.7; very large, r = 0.7–0.9; and nearly perfect, r > 0.9. Alpha level was set at 0.05 for all tests. Data are presented as mean ± 95% confidence interval (t-distribution) upper and lower limits.

## RESULTS

### Participants and motor unit identification and analysis

Twenty-one participants (8 females and 13 males) volunteered for this study (29.2 ± 2.6 years, 72.1 ± 8.8 kg, 174.9 ± 4.9 cm). However, two females and one male did not present motor units with a pulse-to-noise ratio ≥30 dB and were excluded from analysis. Additionally, we were unable to pair the same motor units throughout the different intensities in three participants for *soleus* and four participants for *gastrocnemius medialis*. Thus, their data were excluded from the respective muscle analysis. Therefore, our final sample consisted of 15 participants for *soleus* and 14 participants for *gastrocnemius medialis*.

For the included participants, a mean of 7.5 ± 2.0 motor units were identified for the *soleus* (5.4 ± 1.3 at 10 %; 6.9 ± 1.8 at 20%; and 6.6 ± 2.1 at 30% of MVC) while 10.9 ± 2.6 motor units were identified in *gastrocnemius medialis* (6.8 ± 1.8 at 10%; 9.7 ± 2.5 at 20%; and 8.7 ± 2.3 at 30% of MVC). After tracking the same motor units throughout all contraction intensities, it was possible to identify on average 4.7 ± 1.2 motor units for *soleus* (3.5 ± 2.3 pairs) and 5.4 ± 1.4 for *gastrocnemius medialis* (3.5 ± 2.15 pairs).

The rate-rate correlations between motor unit pairs (first inclusion criteria for paired motor unit analysis) at 10%, 20%, and 30% were 0.85 ± 0.04, 0.93 ± 0.12, and 0.86 ± 0.04 for *soleus* and 0.84 ± 0.04, 0.84 ± 0.03, and 0.84 ± 0.04 for *gastrocnemius medialis*, respectively. The recruitment time differences between test and control units (second inclusion criteria) at 10%, 20%, and 30% were 2.24 ± 0.51, 2.54 ± 0.99, and 2.93 ± 0.80 s for *soleus* and 2.43 ± 0.58, 3.05 ± 0.68, and 2.96 ± 0.51 s for *gastrocnemius medialis*, respectively. There was no statistical difference between 10%, 20%, and 30% conditions for the recruitment time difference between test and control units for *soleus* (F(2-28) = 1.712; p = 0.199; ω^2^ = 0.016) or *gastrocnemius medialis* (F(2-26) = 2.275; p = 0.123; ω^2^ = 0.039). After test unit recruitment, increases in control unit discharge rates (third inclusion criteria) during 10%, 20%, and 30% conditions were 2.73 ± 0.49, 4.24 ± 0.64, and 4.84 ± 0.57 pps for *soleus*, and 2.10 ± 0.48, 3.97 ± 0.70, and 4.60 ± 0.89 for *gastrocnemius medialis*, respectively.

### Delta frequency (ΔF)

The one-way repeated measures ANOVA revealed a large effect for both *soleus* (F _(2-28)_ = 23.696; p < 0.001; ω^2^ = 0.240) and *gastrocnemius medialis* (F _(2-26)_ = 19.780; p < 0.001; ω^2^ = 0.268). In *soleus* (Figure 2A), the post-hoc test identified differences between 10% and 20% (p = 0.001; *d* = 0.991), 10% and 30% (p < 0.001; *d* = 1.773), and between 20% and 30% (p = 0.005; *d* = 0.782). In *gastrocnemius medialis* (Figure 2B), differences between 10% and 20% (p < 0.001; *d* = 1.442) and 10 and 30% (p < 0.001; *d* = 1.470) were identified, however no differences were detected between 20% and 30% (p = 0.917; *d* = 0.030).

### Maximal discharge rate

Maximal *soleus* and *gastrocnemius mediali*s motor unit discharge rates increased with contraction intensity. ANOVA revealed a large effect for both *soleus* (F _(2-28)_ = 34.379; p < 0.001; ω^2^ = 0.363) and *gastrocnemius medialis* (F _(2-26)_ = 89.058; p < 0.001; ω^2^ = 0.453). For *soleus*, differences between 10% and 20% (p < 0.001; *d* = 1.439), 10% and 30% (p < 0.001; *d* = 2.092), and between 20% and 30% (p = 0.017; *d* = 0.653) were identified (Figure 3C), whilst for *gastrocnemius medialis* differences were also identified between 10% and 20% (p < 0.001; *d* = 2.505), 10% and 30% (p < 0.001; *d* = 3.451), and between 20% and 30% (p = 0.002; *d* = 0.946) (Figure 3D).

**Figure 3.**
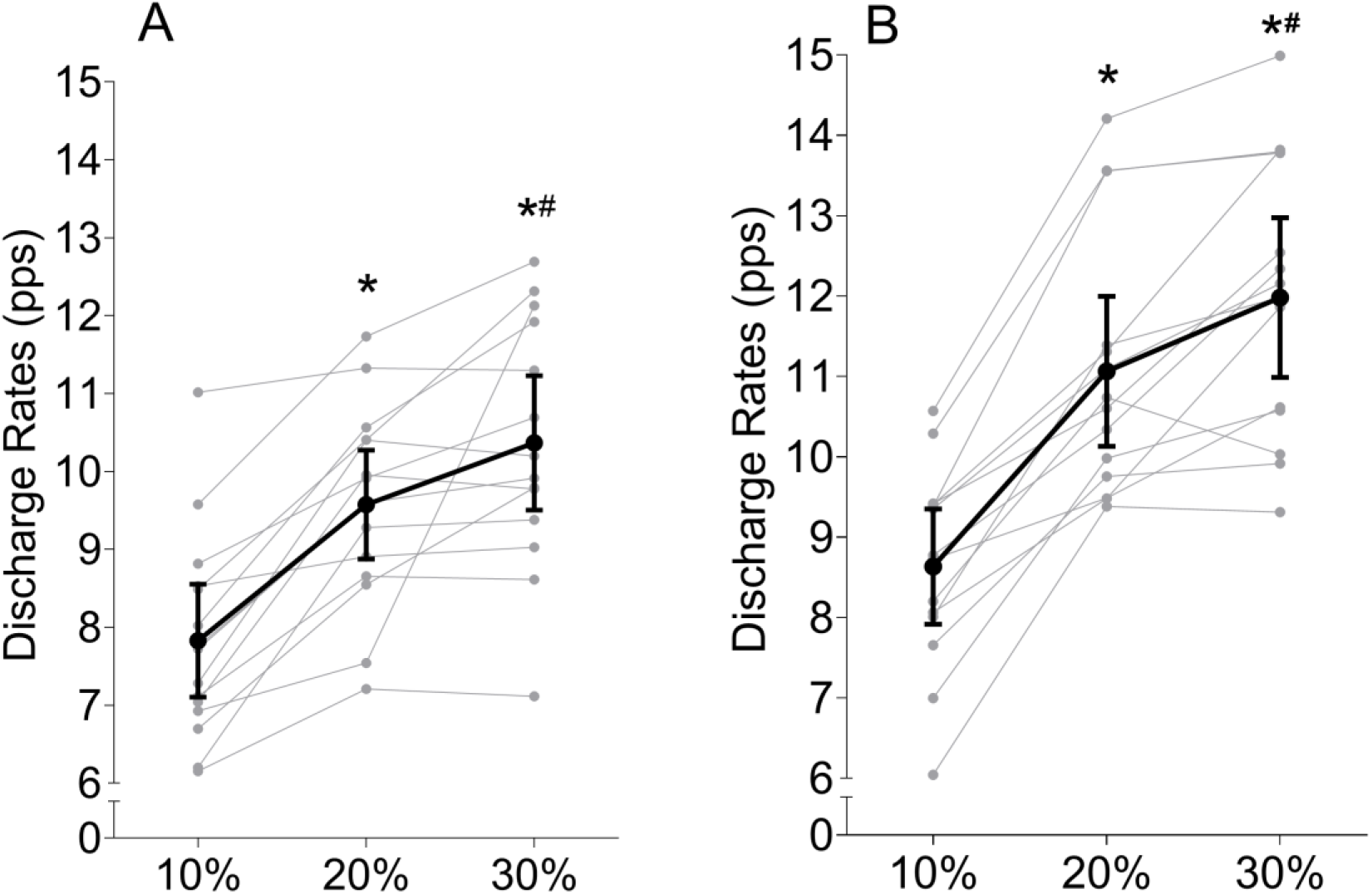
Maximal discharge rate changes across different contraction intensities. Maximal discharge rate of *soleus* (A) and *gastrocnemius medialis* (B) motor units during muscle contractions at 10%, 20% and 30% of maximal voluntary torque. Grey dots and lines show individual results (*soleus*, n = 15; *gastrocnemius medialis*, n = 14) and black dots and lines represent data mean ± 95% confidence interval lower and upper limits. * Different to 10% and # different to 20% conditions.

**Table 1.** Descriptive statistics (mean and mean difference with 95% confidence interval lower and upper limits) for delta frequency (ΔF) and maximal discharge rates for *soleus* and *gastrocnemius medialis* at 10, 20, and 30% of maximal voluntary torque.

| Muscle<br>contraction<br>intensity | *Soleus* |  | *Gastrocnemius medialis* |  |
| --- | --- | --- | --- | --- |
| | $\Delta F$ (pps) | Maximal<br>discharge rate<br>(pps) | $\Delta F$ (pps) | Maximal<br>discharge rate<br>(pps) |

| Mean |  |  |  |  |
| --- | --- | --- | --- | --- |
| 10 % | 2.05<br>(1.66 – 2.43) | 7.83<br>(7.10 – 8.55) | 1.79<br>(1.28 – 2.30) | 8.63<br>(7.92 – 9.35) |
| 20 % | 2.63*<br>(2.28 – 2.99) | 9.58*<br>(8.88 – 10.28) | 3.10*<br>(2.52 – 3.69) | 11.06*<br>(10.13 – 12.00) |
| 30 % | 3.09*#<br>(2.61 – 3.58) | 10.37*#<br>(9.50 – 11.23) | 3.13*<br>(2.48 – 3.78) | 11.98*#<br>(10.99 – 12.97) |
| Mean difference |  |  |  |  |
| 20 – 10 % | 0.58<br>(0.33 – 0.84) | 1.75<br>(1.23 – 2.26) | 1.32<br>(0.83 – 1.81) | 2.43<br>(1.82 – 3.03) |
| 30 – 20 % | 0.46<br>(0.14 – 0.78) | 0.80<br>(0.08 – 1.51) | 0.03<br>(-0.34 – 0.39) | 0.92<br>(0.36 – 1.48) |

\* Different to 10% and # different to 20% conditions.

### Delta frequency (ΔF) vs Maximal discharge rate

The repeated-measures correlation between maximal discharge rates and ΔFs across the contractions intensities was *large* for *soleus* (r = 0.640; 95% confidence interval = 0.358 - 0.815; p < 0.001; Figure 4A) and *very large* for gastrocnemius medialis (r = 0.773; 95% confidence interval = 0.557 – 0.819; p < 0.001; Figure 4B).

**Figure 4.**
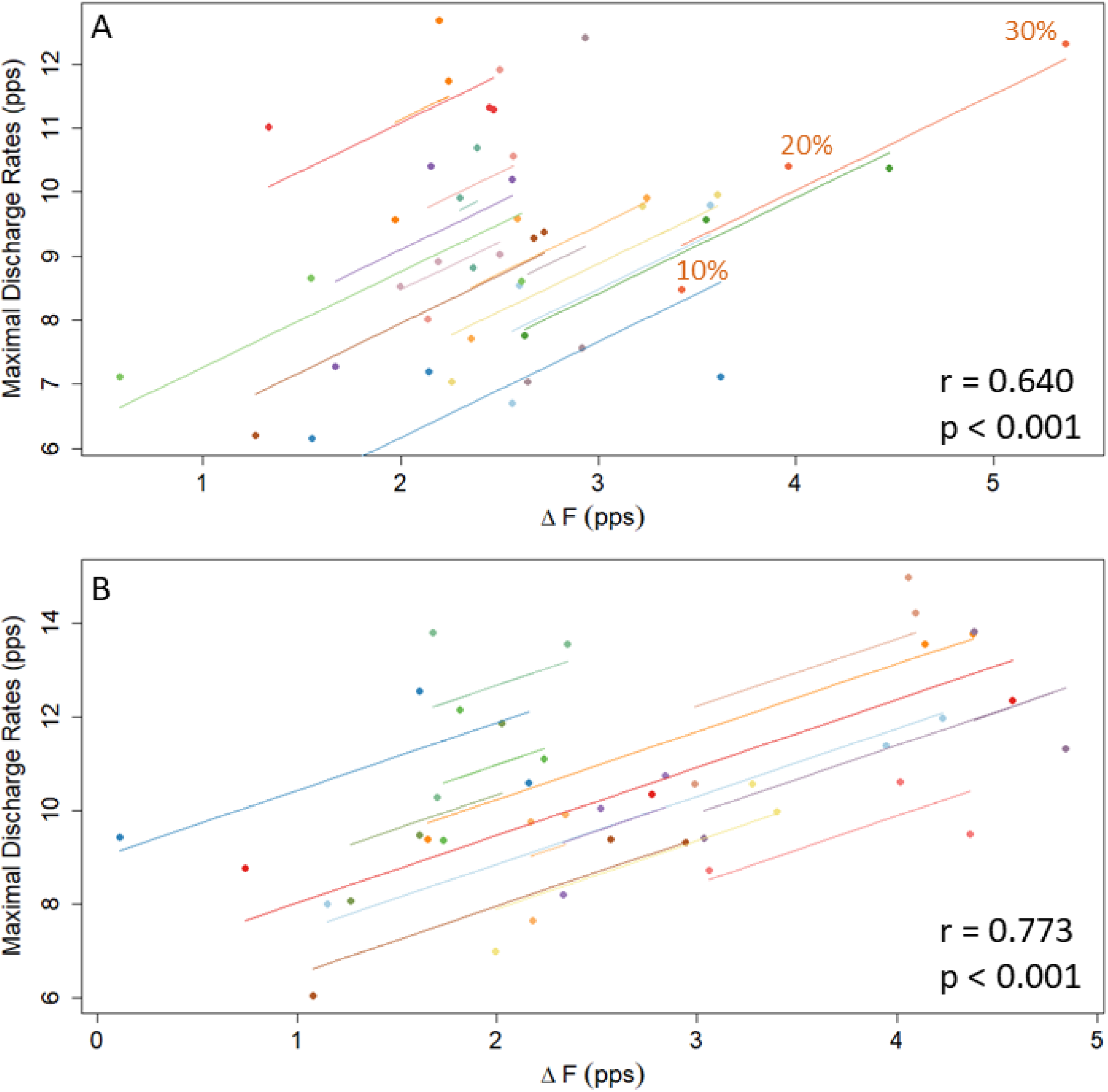
Repeated-measures correlations plot illustrating the association between the maximal discharge rates and ΔF across different contractions intensities. Panel A depicts results for *soleus* and panel B the results for *gastrocnemius medialis*. Separate parallel lines are fitted to the data from each participant across contractions intensities (10%, 20%, and 30% of MVC) and are represented by different colours.

**Figure 5.**
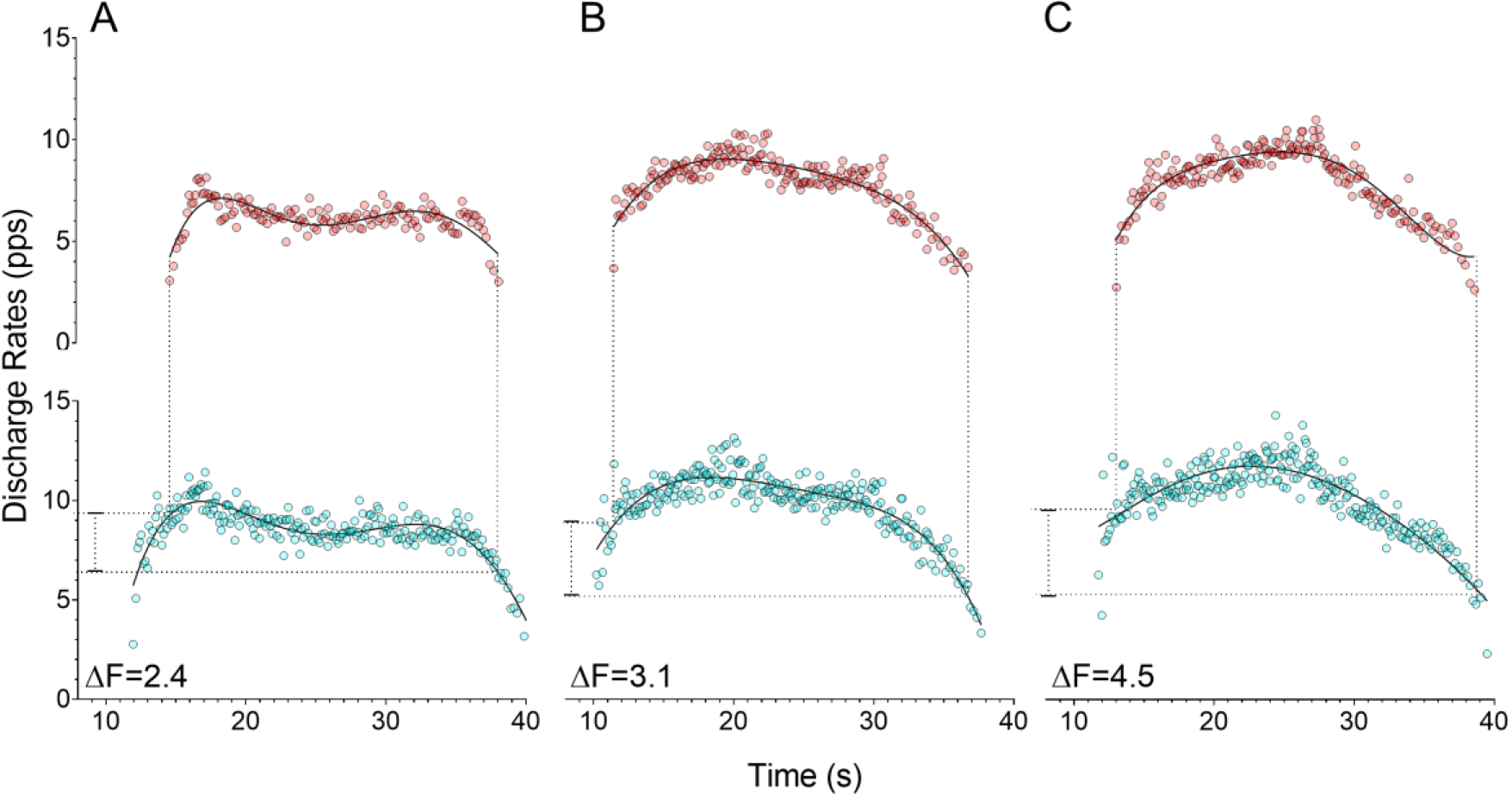
Delta frequency (ΔF) calculation. Paired-motor unit analysis for the ΔF calculation from the same *soleus* motor units from one participant. Test units (top graphs - red motor units) and control units (bottom graphs – blue motor units) during the ramped isometric contractions at 10% (A), 20% (B) and 30% (C) of maximal voluntary torque. The black lines represent the fifth order polynomial fit used for the ΔF calculation.

## DISCUSSION

The present study was designed to determine whether PIC amplitudes, estimated using paired-motor unit ΔFs analysis, increase according to the level of voluntary motor drive in human *soleus* and *gastrocnemius medialis* motor neurones. As the motor units obtained at the lowest contraction intensity (10% MVC) were then tracked across conditions, the study focussed only on low-recruitment threshold motor units. Our results showed a significant increase in ΔFs motor units from 10% to 20% (Δ = 0.6 pps) and 20% to 30% (Δ = 0.5 pps) of maximal voluntary contraction (MVC) in *soleus* and from 10% to 20% (Δ = 1.3 pps) of MVC in gastrocnemius medialis. Increases in ΔF were also positively correlated with maximal discharge rates (output) for both *soleus* (r = 0.640) and gastrocnemius medialis (r = 0.773). These findings support the hypothesis that increases in lower-threshold motor units’ PIC amplitude might act as a variable gain control in humans, increasing the gain as force production increases from 10% to 30% of MVC. However, they are also suggestive of gain adjustments differing between synergistic muscles, as both muscles showed different ΔF increases despite similar firing rate increases. These results not only indicate that PICs may play an important role in varying input-output gain to control muscle force production, but that between-muscle or -person ΔF comparisons using the paired motor unit technique would need to be done with the muscles achieving the same relative level of force output (or activation).

As evidenced by our results, ΔF appears to increase in proportion to motor output, and presumably the level of descending drive, in human *soleus* and *gastrocnemius medialis* muscles, as the increases in ΔF and maximal discharge rate (i.e., indirect measure of muscle activation) across contraction intensities were correlated. These results are consistent with previous research in animal models (Powers, Nardelli and Cope, 2008; Huh *et al*., 2017) as well as computational modelling studies (Heckman and Binder, 1991; Heckman, 1994; Randall K. Powers and Heckman, 2015). In animals, this increase in input-output relationship has been shown to be promoted by the presence of serotonin and noradrenaline in the spinal cord. For example, exogenous application of serotonin in the spinal cord increases locomotion speed (i.e., motor output) in zebrafish larvae and in newborn rats (Cazalets, Sqalli-Houssaini and Clarac, 1992; Brustein *et al*., 2003). In humans, selective pharmacological manipulation of serotonin levels directly affects motor neurone excitability (Wei *et al*., 2014; Kavanagh, McFarland and Taylor, 2019). Circulating serotonin and noradrenaline levels are increased during higher-intensity physical exercise (higher motor output) compared to a resting state (lower or no motor output) (Chaouloff, 1989; Ramel, Wagner and Elmadfa, 2004; Udina *et al*., 2010). Thus, it is reasonable to speculate that the increased ΔFs observed at higher contraction intensities in the present study could result from greater serotonergic and/or noradrenergic drive. Therefore, we suggest that the contraction intensity-dependent increase in PIC amplitudes in *soleus* and *gastrocnemius medialis*, indicated by the increases in ΔFs, may result from a greater serotonergic and/or noradrenergic drive from brainstem nuclei as voluntary drive is increased (Johnson and Heckman, 2014; Binder, Powers and Heckman, 2020). Our results corroborate the gain control system theory developed in animal and computer modelling experiments (Johnson and Heckman, 2014). Higher ΔFs accompanying both the greater torque output and maximal discharge rates indicates that this gain control system is also observed in human motor neurones.

An interesting finding was that the ΔF increase per increment in total muscle force differed between the muscles, despite them showing similar mean overall ΔF increases from 10-30% of MVC (1.1 pps for *soleus* and 1.3 pps for *gastrocnemius medialis). Soleus* ΔF increased from 10 to 20% of MVC (from 2.05 to 2.63 pps) and then increased further to 30% (2.63 to 3.09 pps) whereas a greater increase was observed from 10 to 20% MVC (1.79 to 3.10 pps) in *gastrocnemius medialis* without further change from 20% to 30% MVC (3.10 to 3.13 pps). One interpretation of this result is that *soleus* ΔF increased at a smaller but more consistent rate than in *gastrocnemius medialis*, which could indicate an early saturation of the contraction intensity-PIC amplification relationship in *gastrocnemius medialis* motor neurones. That is, the identified *gastrocnemius medialis* motor units reach their maximal amplification level at a lower level of force, and presumably lower level of voluntary drive, than *soleus*. Differences in PIC amplitudes between muscles and contraction intensities could be related to the intrinsic properties of the motor neurones themselves. For example, PICs appear to last longer in the slow-type motor units’ neurones (Heckman *et al*., 2008), which are more abundant in postural muscles such as *soleus* (~80-70%) than muscle such as *gastrocnemius medialis* (~55-60%) (Gollnick *et al*., 1974; Harridge *et al*., 1996; Houmard *et al*., 1998).

Interestingly, our results are contrary to two recent studies that compared ΔFs obtained from the same three contraction intensities in humans (i.e., 10, 20, and 30% of MVC) (Afsharipour *et al*., 2020; Kim *et al*., 2020). Kim et al., (Kim *et al*., 2020) tested *soleus* and *tibialis anterior*, while Afsharipour et al., (Afsharipour *et al*., 2020) tested *tibialis anterior*, and none of them reported significant differences for ΔFs among contraction intensities. A few methodological differences between their studies and ours might explain the divergence observed in our results. First, they (Afsharipour *et al*., 2020; Kim *et al*., 2020) adopted distinct torque rise among contraction intensities (i.e., 10-s ramp up and 10-s ramp down for all intensities, resulting in 1%, 2%, and 3% torque rise per sec for 10%, 20%, and 30% of MVC, respectively). Quicker rates of torque rise can reduce ΔFs due to spike-threshold accommodation (Randall K. Powers and Heckman, 2015), which may have underestimated the ΔFs when contraction intensity was increased. Second, Kim et al., (Kim *et al*., 2020) and Afsharipour et al., (Afsharipour *et al*., 2020) did not track the same motor units across contractions intensities. As previously reported (Hassan *et al*., 2020), there is a large between-motor unit variability in ΔFs, which might have reduced the sensitivity of the variable to detect differences among contractions intensities. Third, both studies compared ΔFs obtained from different populations of motor units. Specifically, while the lower intensity condition (10% MVC) included only motor units recruited from 0-10% of maximal torque, higher intensities (20 and 30% MVC) included motor units recruited from 0-20% and 0-30% of maximal torque, respectively. Also, the proportion of lower-threshold motor units decomposed by the algorithm seems to be reduced in higher contractions intensities (Hassan *et al*., 2019). Therefore, the authors might have compared mostly lower-vs higher-threshold motor units for lower vs higher contraction intensities, respectively. The use of motor units of different recruitment threshold for different contraction intensities limit the interpretation of the independent effect of contraction intensity on ΔF values in these studies.

One strength of the present study was that both the rate of torque rise and decline during ramp-shaped contractions and the contraction durations were identical between conditions. This was necessary to control for potential influences of spike frequency adaptation and spike-threshold accommodation on PIC amplitude (Vandenberk and Kalmar, 2014; Randall K. Powers and Heckman, 2015). However, torque path traces were different, and it is possible that different strategies might have been used to accomplish these tasks. Another strength of the present study was that the same motor units were tracked across different contraction intensities, which allows the comparison of motor units receiving the same synaptic input. Also, a previous work from our group presented a very good reliability for ΔFs in a repeated-measures design when the motor units were compared across different contractions (Trajano *et al*., 2020). However, this choice also reduced the total number of motor units available for study, limiting our analysis to lower-recruitment threshold motor units which initiated firing already within the lowest-intensity contraction (i.e. 10% of MVC). Thus, it remains to be determined whether these findings are obtained when higher-threshold units are studied. Based on our findings and considering the study design limitations, caution should be observed when extrapolating our findings to higher-recruitment threshold motor units and different muscles.

In conclusion, the present data suggest that estimates of PIC amplitudes (ΔFs) in low-threshold motor units increase with the level of voluntary descending drive in human *soleus* and *gastrocnemius medialis* muscles. *Soleus* PIC amplitude increased consistently across contractions from 10 to 30% of MVC, while for *gastrocnemius medialis* PIC amplitude increased more prominently from 10 to 20% of MVC but did not increase further during the 30% MVC contraction. These results were associated to increases in maximal motor unit discharge rate with increasing contraction intensity from 10 to 30% of MVC in both *soleus* and *gastrocnemius medialis*. It appears that increases in muscle activation with increasing force occur through both an increase in voluntary descending drive and concomitant increase in PIC amplitudes, providing both cortical- and spinal-based mechanisms of muscle recruitment. Also, the data indicate that comparisons between studies that adopt different contraction strengths may not be reasonable when using the paired MU technique, and that between-muscle comparisons may only be performed if the descending drive (or muscle force) is of the same relative level between the muscles. Our findings indicate only the behaviours of low-recruitment threshold motor units in *soleus* and *gastrocnemius medialis* muscles, so future research could investigate changes PIC amplitudes at higher contraction intensities (e.g., 40 – 100% MVC), and thus in higher threshold motor units, and in different muscles.

